# c-MAF maintains the transcriptional program of enterocyte zonation and the balance of absorptive/intestinal secretory cell types

**DOI:** 10.1101/2021.12.03.471081

**Authors:** Alejandra González-Loyola, Tania Wyss, Olivia Munoz, Borja Prat-Luri, Mauro Delorenzi, Gregory Verdeil, Tatiana V. Petrova

## Abstract

Small intestinal villi are structural and functional units uniquely adapted to the nutrient absorption in higher vertebrates. Villus enterocytes are organized in spatially resolved “zones” dedicated to specialized tasks such anti-bacterial protection, and absorption of amino-acids, carbohydrates and lipids. The molecular mechanisms specifying villus zonation are incompletely understood. We report that inactivation of transcription factor c-MAF, highly expressed in mature lower and mid-villus enterocytes, perturbed the entire villus zonation program, by increasing the expression of regulators of carbohydrate and bile acid metabolism and transport, while suppressing genes related to amino acid and lipid absorption. *Maf* inactivation under homeostatic conditions expanded tuft cells and led to compensatory gut lengthening, preventing body weight loss. However, delayed enterocyte maturation in the absence of *Maf* impaired body weight recovery after acute intestinal injury, resulting in reduced survival. Our results identify c-MAF as a novel regulator of small intestinal villus zonation program, while highlighting the importance of coordination between stem/progenitor and differentiation programs for intestinal regeneration.

**Summary:** c-MAF is expressed in differentiated enterocytes. c-MAF loss alters enterocyte zonation leading to a compensatory gut remodelling and tuft cell expansion. Upon acute intestinal injury mice deficient for c-MAF cannot recover due to lack of nutrient transport and compensatory lengthening.

## Introduction

The intestinal epithelium is one of the most dynamic structures in our body that continuously regenerates to replace damaged cells and ensure proper intestinal functions, such as absorption of nutrients and water. In mice, this constant regeneration replenishes the entire epithelium every 5-7 days and is driven by intestinal stem cells residing in the crypts. Intestinal stem cells give rise to all differentiated epithelial populations: enterocytes, responsible for nutrient absorption, Paneth cells, which synthesize and secrete antimicrobial peptides and proteins, mucus-producing goblet cells, tuft cells and enteroendocrine cells, which secrete a variety of hormones (Barker, 2014; Sato et al., 2009). Perturbed intestinal epithelial regeneration is also a frequent and severe side-effect of anti-cancer therapies (Yu, 2013). Thus, understanding the molecular mechanisms that drive intestinal epithelial homeostasis at steady-state and after exposure to anti-cancer therapies could provide novel therapeutic tools.

Much attention has been paid to the characterization of the intestinal stem and progenitor cell compartments (Barker et al., 2007; Clevers, 2013; Guiu et al., 2019). However, the recent discovery of the functional compartmentalization of enterocytes along the intestinal villus axis has also highlighted the heterogeneity and complexity of the differentiated epithelial cells (Moor et al., 2018). This zonation extends to the nutrient absorption pathways as mid-villus enterocytes absorb aminoacids and carbohydrates, while villus tip enterocytes take up lipids and produce chylomicrons (Moor et al., 2018). Gut lumen extracellular ATP is a signal of danger capable of activating the intestinal immune system (Trautmann, 2009). At the villus tip, purine metabolites, such as adenosine and inosine, promote an anti-inflammatory program by decreasing leukocyte infiltration and inflammatory cytokine release from activated macrophages (Mabley et al., 2003). Furthermore, villus tip enterocytes express 5’ nucleotidase (NT5E) that transforms AMP to adenosine and *Nt5e* knockout mice develop autoimmune reactions (Blume et al., 2012) and cannot recover from induced colitis (Bynoe et al., 2012). Therefore, villus tip enterocytes are necessary to prevent exaggerated immune responses to the gut environment.

The highly conserved MAF family of transcription factors harbor a DNA binding domain that allows binding to specific DNA sequences called MAREs (Maf-recognition elements) (Kerppola and Curran, 1994). Within the family, c-MAF regulates transcriptional programs involved in differentiation processes such as lens fiber and tubular renal cell differentiation (Imaki et al., 2004; Kawauchi et al., 1999) and chondrocyte development (Hong et al., 2011). In the pancreas c-MAF activates glucagon gene expression (Kataoka et al., 2004) and in fetal liver it regulates erythropoiesis (Kusakabe et al., 2011). Since its identification as a regulator of Th2 immune cells (Ho et al., 1996), knowledge about its role in immune cell homeostasis has exponentially grown. c-MAF is expressed in innate immune cells (Parker et al., 2020; Pokrovskii et al., 2019), B lymphocytes (Liu et al., 2018) and T cell subsets such as Type 1 regulatory cells (Pot et al., 2009), T helper 17 (Hiramatsu et al., 2010), CD8 T cells (Giordano et al., 2015) and T follicular helper cells (Andris et al., 2017), but also in type 2 macrophages (Daassi et al., 2016). Recently, it was reported that c-MAF regulates the balance between TH17 and Treg numbers in the gut and thus maintains intestinal homeostasis (Imbratta et al., 2019). Here, we report a novel role of c-MAF in murine differentiated enterocytes of the small intestine.

## Results and Discussion

### c-MAF is expressed in small intestinal enterocytes in response to BMP signalling

A survey of different murine tissues revealed high *Maf* expression in mouse eye, spleen, fat, ovary, muscle, diaphragm, kidney, and small intestine (**Fig.1a**). To identify the cell types expressing *Maf* in the gut we stained small intestinal sections for c-MAF protein, and we analysed *Maf* mRNA levels in sorted EPCAM1+/CD44+ progenitor cells and mature EPCAM1+/CD44-enterocytes (Snippert et al., 2009; Zeilstra et al., 2008). *Maf* mRNA and protein were highly expressed in mature villus CD44neg enterocytes and absent in CD44+ progenitor/stem cells or secretory cell types (**Fig.1b-e, Fig.S1a, b**). Expression of *Maf* in mature enterocytes but no other intestinal epithelial lineages was further confirmed in a single cell RNA sequencing (scRNAseq) dataset of mouse small intestinal epithelial cells (**Fig.1e, Fig.S1c-d** (Yan et al., 2017). Along the gut, c-MAF was highly expressed in small intestinal villous enterocytes, while we observed only low levels of c-MAF in colon epithelial cells (**Fig.S2a**).

**Figure 1.**
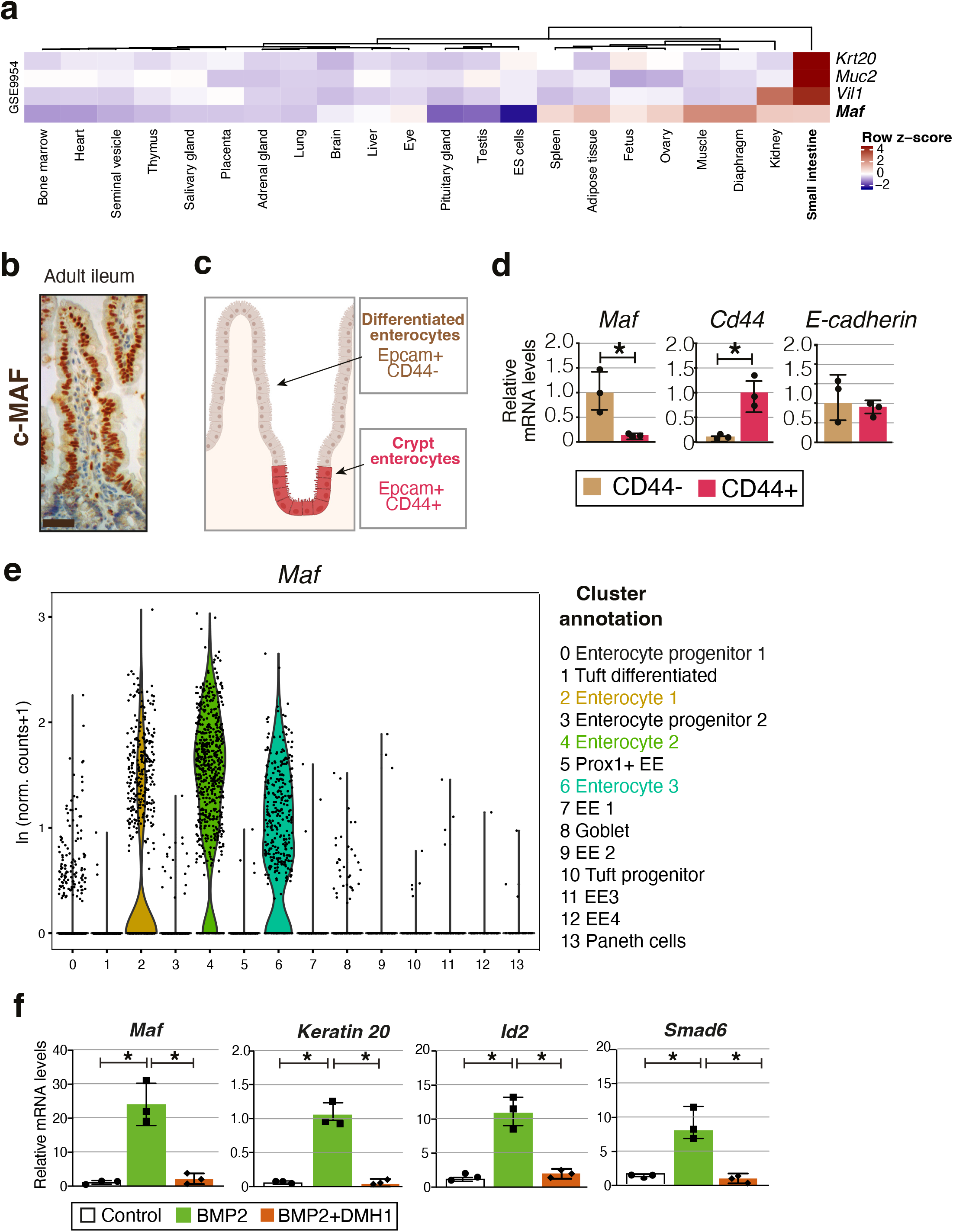
c-MAF is expressed in small intestinal enterocytes and induced upon differentiation. **a**. Expression profile of *Maf* and intestinal epithelial genes *Krt20, Muc2, Vil1* in mouse tissues (GSE9954). A high z-score (dark red) indicates higher expression of the genes in specific tissues. **b**. Staining for c-MAF (brown) in adult mouse ileum. Scale bar, 25 μm. **c**. Schematic representation of the location of Epcam+Cd44+ and Epcam+Cd44-cells along the villus and sorted from WT small intestine. **d**. *Maf* is expressed in Epcam+Cd44-cells. RT-qPCR analysis of the indicated genes. Mean ± SD. *, p < 0.05; n = 3 WT, n = 3. **e**. *Maf* is expressed in mature enterocytes. Expression level per cell (ln[normalized counts + 1]) of *Maf* overlaid on scRNAseq dataset of Yan et al., 2018. **f**. BMP2 induces *Maf*. WT small intestinal organoids were treated for 24 hours with 100 ng/ml recombinant BMP2 or BMP2+ 3μm DMH1. Relative expression levels of *Maf, Krt20, Id2, Smad6* are shown as mean ± SD. *, p < 0.001; n = 3 different organoid preparations.

During embryonic development, c-MAF was absent in E12.5 gut and sporadic c-MAF positive gut epithelial cells were observed at E17.5 (**Fig.S2b-c**). In contrast, all villus enterocytes expressed c-MAF at P4 and thereafter (**Fig.1b, Fig.S2a, d**). Postnatal microbial colonization plays an important role in the development and shaping of intestinal immune cells (Gensollen et al., 2016), however epithelial c-MAF levels were unchanged in mice treated with broad-spectrum antibiotics (data not shown), indicating that its expression after E12.5 is part of the intrinsic epithelial maturation program.

Gradients of WNT and BMP signalling maintain the balance between proliferating crypts and differentiated villous epithelial cells (Clevers, 2006). Given high expression of c-MAF in differentiated intestinal epithelial cell *in vivo*, we isolated small intestinal organoids and treated them with BMP2. As expected, BMP2 treatment induced expression of its target genes *Id2* and *Smad6* (Hollnagel et al., 1999; Takase et al., 1998), as well as the marker of differentiated intestinal epithelium *Krt20* (Moll et al., 1990). BMP2 also strongly promoted the expression of *Maf*, and this induction was abolished when BMP2 signalling was inhibited by the selective BMP inhibitor DMH-1 (**Fig.1f)**. Collectively, our data establish that c-MAF is expressed in mature small intestinal enterocytes *in vivo* and that its expression is induced upon differentiation *in vitro*.

### Loss of *Maf* perturbs the transcriptional program of enterocyte zonation

To understand the role of c-MAF in enterocytes, we generated *Maf*^fl/fl^*;Vil-*CreERT2 *(Maf*^IECKO^) mice in which administration of tamoxifen leads to depletion of *Maf* in intestinal epithelial cells (**Fig.2a**). Despite the profound loss of c-MAF protein and mRNA, and the absence of compensatory increase in other related MAF family members (**Fig. 2a,b Fig.S3a** (Kataoka et al.2007), we did not observe changes in the general enterocyte maturation status as determined by staining for KRT20 (**Fig.2c**(Chan et al., 2009) or in epithelial cell proliferation or turnover as determined by the analysis of EdU^+^ epithelial cells distribution 1h and 48h after EdU administration (**Fig.S3b**). Furthermore, the body weight of *Maf*^IECKO^ mice was similar to that of WT littermates (**Fig.2d**), indicating globally unaltered nutrient absorption capacity.

**Figure 2.**
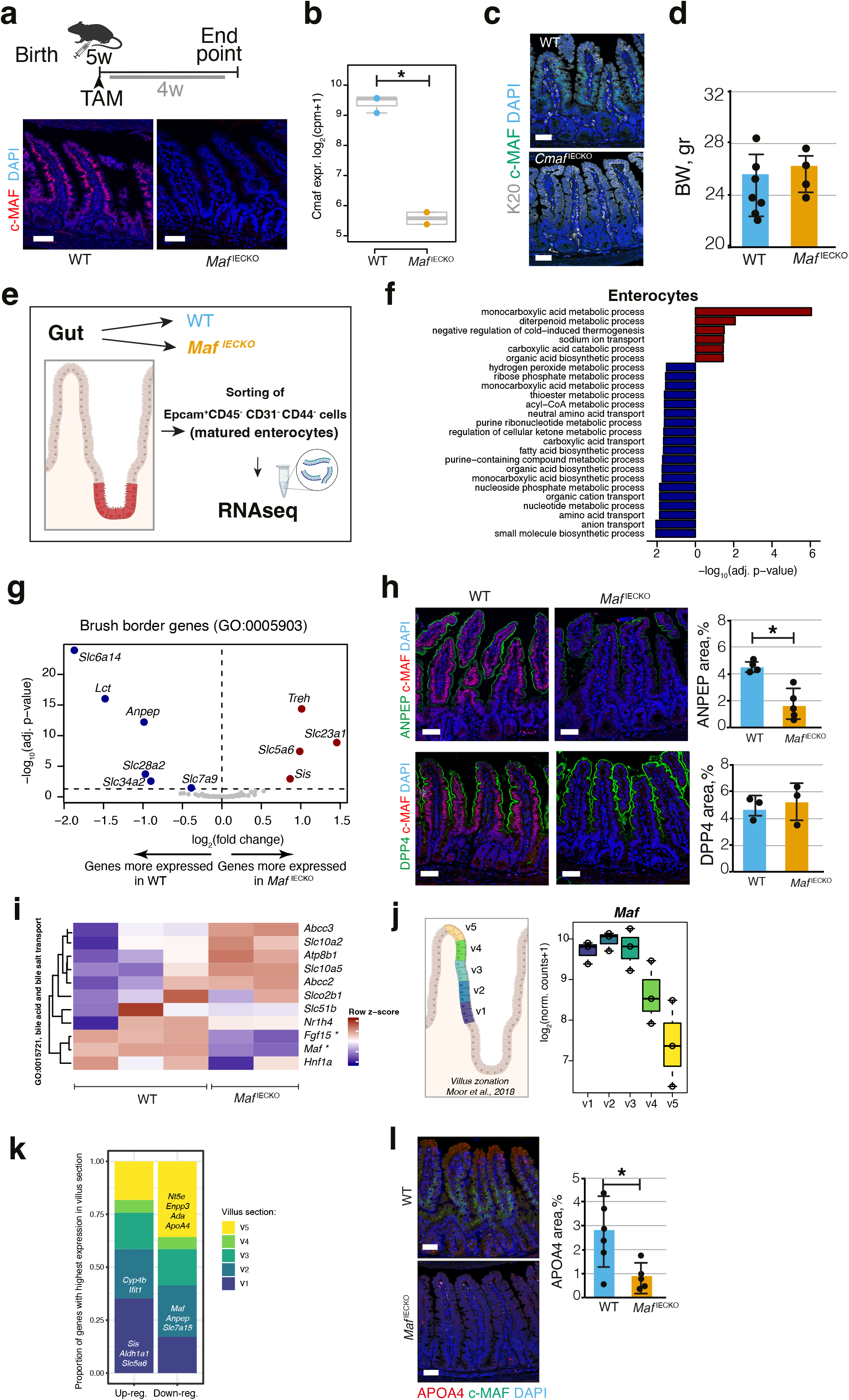
Loss of *Maf* in adult enterocytes alters their transcriptional zonation program. **a**. *Maf* depletion in *Maf*^IECKO^ mice. Staining for c-MAF (red) and DAPI (blue). c-MAF levels were analyzed in WT and *Maf*^IECKO^ mice 1 month after the initiation of depletion. n=10 WT, n =11 *Maf*^IECKO^ from 3 different cohorts. Scale bar, 50μm. **b**. *Maf* levels are significantly reduced in *Maf*^IECKO^ *n* = 2 WT; *n* = 3 *Maf*^IECKO^ *, p < 0.01. **c**. *Maf* depletion does not change overall enterocyte differentiation. Staining for keratin 20 (K20, white), c-MAF (green) and DAPI (blue). Scale bar, 50μm. **d**. *Maf* depletion does not affect mouse weight. Mean ± SD. *n* = 7 WT; *n* = 4 *Maf*^IECKO^. n.s.: not significant. **e**. Sorting strategy for isolation of mature enterocytes from WT and *Maf*^IECKO^ mice for RNAseq. **f**. Signalling pathways enriched in *Maf*^IECKO^ vs WT mouse mature enterocytes. Bar plots of GO gene sets significantly over-represented (red bars) or under-represented (blue bars) in *Maf*^IECKO^ vs WT mice after 1 week of *Maf* depletion. n=3 WT, n =2 *Maf*^IECKO^. NES, normalized enrichment score; adjusted *p*-value = 0.05. **g**. Volcano plot of differentially expressed brush border genes (GO: 0005903) in *Maf*^IECKO^ mice. **h**. ANPEP but not DPP4 is reduced upon *Maf* depletion. Staining for ANPEP or DPP4 (green), c-MAF (red) and DAPI (blue). Scale bar, 50μm. Percentage of area per tissue in WT and *Maf*^IECKO^ mice, mean ± SD. *, p < 0.05. n=3 WT; n=3 *Maf*^IECKO^. **i**. Heatmap of bile acid regulators, *Maf* and *Fgf15* were significantly different in *Maf*^IECKO^ *versus* WT mice. **j**. *Maf* expression is zonated along small intestinal villus, as observed along villus zones described in Moor et al., 2018. **k**. Proportion of genes differentially expressed upon *Maf* loss with highest expression in the 5 different villus zones described in Moor et al., 2018. Most genes induced in the absence of *Maf* have their highest expression level in proximal zone v1, while most decreased genes in the absence of *Maf* are highly expressed in villus tip zone v5. Examples of transcripts characteristic of each zone are indicated. **l**. Marker of v4 and v5 zones APOA4 is reduced in *Maf*^IECKO^ mice. Staining for c-MAF (green), APOA4 (red) and DAPI (blue). Scale bar, 50μm. Percentage of area in WT and *Maf*^IECKO^ mice, mean ± SD. *, p < 0.05. n=6 WT; n=5 *Maf*^IECKO^.

To characterize the impact of *Maf* loss on the enterocyte transcriptional program we performed RNA sequencing (RNAseq) analysis of mature CD44neg small intestinal enterocytes sorted from WT or *Maf*^IECKO^ mice. Loss of *Maf* resulted in 106 differentially upregulated genes and 77 downregulated genes (adjusted *p*-value < 0.05, mean expression level > 500 normalized counts, **Table S1**). Unbiased analysis of signaling pathways using gene set enrichment analysis (GSEA, (Subramanian et al., 2005) indicated defective metabolic nutrient pathways for amino and fatty acids, while pathways associated with carbohydrate metabolism were increased (**Fig.2e,f Table S1**). The brush border is a hallmark of mature enterocytes essential for terminal digestion and absorption of nutrients (Carboni et al., 1987). Therefore, we analyzed the expression of brush border transcripts in *Maf*^IECKO^ mice. We found that in the absence of *Maf*, transcripts involved in carbohydrate metabolism such as treholase (*Treh*) or sucrose isomaltase (*Sis*) (Hauri et al., 1979) were enhanced, whereas the aminoacid transporters *Slc7a9, Slc6a14* (Karunakaran et al., 2011), the aminopeptidase *Anpep* needed for the digestion of dipeptides (Luan and Xu, 2007) or the purine transporter *Slc28a2* (Young et al., 2013) were diminished (**Fig.2g**). At the same time the expression of other genes associated with enterocyte differentiation such as *Krt20, Fabp1* or *Dpp4* were not affected (**Table S1** and (Bass, 1988; Darmoul et al., 1992). We confirmed the RNAseq data analysis by immunohistochemistry for ANPEP, which was downregulated in *Maf*^IECKO^ mice, whereas the dipeptidyl peptidase 4 (DPP4) remained unchanged (**Fig.2h**).

In addition to changes in the nutrient transporters, the atypical fibroblast growth factor 15 (*Fgf15*) that binds hepatocyte FGF receptor 4 to repress bile acid synthesis (Inagaki et al., 2005) was strongly reduced in *Maf-*depleted enterocytes (**Fig.S3c)**. Consistently, transcriptome analysis showed increased expression of the main bile acid transporter *Slc10a2* which is expressed similar to *Maf* in lower and mid villi (**Fig.2, i, j** and data not shown). Collectively, these results indicate that loss of *Maf* does not impact overall differentiation status of enterocytes but rather fine tunes the transcriptional programs for nutrient and bile acid absorption.

Mature enterocytes are organized in transcriptionally and functionally specialized zones along the villus axis (Moor et al., 2018). Therefore, we analyzed the expression of *Maf* along the villus based on the intestinal zonation pattern described by Moor et al., 2018. As observed by immunohistochemistry (**Fig.2a**), the highest levels of *Maf* were present in *the* lower and mid villus (v1, v2, v3), while *Maf* expression declined towards the villus tip in zones v4 and v5 (**Fig.2j**). Next, we deconvoluted the zonation pattern of genes differentially expressed in *Maf*^*iECKO*^ vs wild type (WT) enterocytes by determining whether they were most highly expressed in the lower, mid or tip of villi in the data of Moor et al., 2018. Significant transcriptional changes were observed in the v2 zone, where *Maf* is highly expressed, and where main intestinal transporters including carbohydrate and aminoacid transporters such as *Slc16a10, Slc43a2, Slc7a7* (Bröer, 2008) and peptidases such as *Anpep* are mainly located (**Table S1**).

Surprisingly, we found that loss of *Maf* significantly decreased many transcripts located in zone 5 at the tip of the villus, where *Maf* levels are low. Notably, we observed downregulation of *Apoa4*, needed for assembly of chylomicrons (Kohan et al., 2015) and multiple purine catabolism genes including *Nt5e, Ada or Npp3, Slc28a2* (Huber-Ruano et al., 2010; Moor et al., 2018; Robson et al., 2006) (**Fig.2k, Table S1**). This result was validated by immunohistochemistry for APOA4 which was significantly diminished at the tip of the villi of *Maf*^*iECKO*^ mice (**Fig.2l**). These data indicate that in the absence of *Maf* enterocytes migrating toward the villus tip are unable to undergo further specialization.

To study the evolutionary conservation of c-MAF function in intestinal epithelium we analysed the expression of human *c-MAF* or its zebrafish ortholog *mafa* scRNA seq data sets. *c-MAF* was highly expressed in differentiated human ileal enterocytes, which also expressed high levels of ANPEP and APOA4, but not in progenitor/stem cells or secretory lineages (**Fig. S3d**). Among zebrafish intestinal populations, *mafa* was strongly produced by enterocytes 3 and 4, involved in lipid and carbohydrate absorption (Willms et al., 2021), while secretory or progenitor cells were negative. Of interest, lysosome-rich enterocytes involved in protein catabolism had the highest levels of *mafa* (**Fig. S3e**, (Park et al., 2019). These results are consistent with conservation of *Maf* function in the regulation of intestinal nutrient absorption transcriptional programs in all vertebrates.

Taken together, our results indicate that c-MAF is dispensable for transition from progenitor to differentiated enterocytes. However, it is important to maintain a balanced zonated transcriptional program for purine catabolism and nutrient and bile acid transport in mature enterocytes along intestinal villi.

### Loss of *Maf* leads to compensatory gut remodeling and expansion of tuft cells at steady state

We were puzzled that despite prominent transcriptional changes in the intestinal epithelium, *Maf*^IECKO^ mouse weight did not differ from those of the WT littermates. We therefore analyzed the length of the intestine and the size of crypt and villous compartments in *Maf* deficient and WT mice. We found that small intestine but not colon of *Maf*^IECKO^ mice was significantly longer as compared to that of WT mice (**Fig.3a, Fig.S4a**), indicating a compensatory enlargement of the absorptive surface. In agreement with this observation, we found that small intestinal crypts area and villus length were significantly greater in *Maf*^IECKO^ animals (**Fig.3b**). Of note, *Maf* inactivation did not increase crypt fission events (**Fig.S4b**), which represents another way of increasing the intestinal surface (Dehmer et al., 2011).

**Figure 3.**
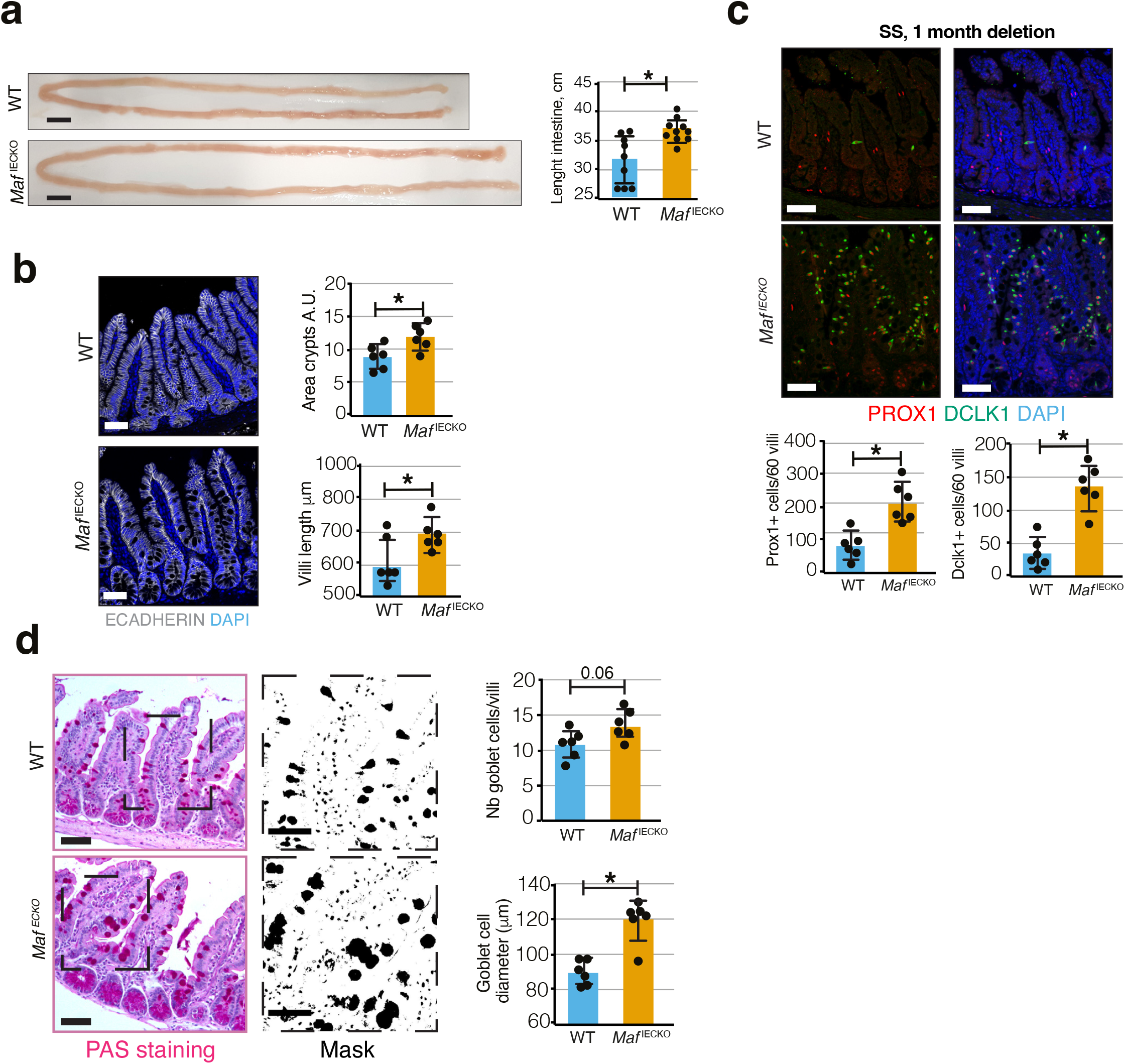
Loss of *Maf* in adult intestinal epithelium leads to gut remodelling and tuft cell expansion. **a**. Increased length of small intestine in *Maf*^IECKO^ mice. Scale bar, 1 cm. *, p < 0.05. n = 9 WT, n = 10 *Maf*^IECKO^. **b**. Increased villi length and crypt area upon loss of *Maf*. Staining for E-cadherin (white) and DAPI (blue). Quantification of crypt area and villi length of small intestine. *, p < 0.05. n = 6 WT, n = 6 *Maf*^IECKO^. **c**. Tuft cell frequency is increased upon *Maf* depletion. Staining for PROX1 (red), DLCK1 (green) and DAPI (blue). Number of PROX1^+^ and DCLK1+ cells in 60 villi. (mean ± SD). *, p < 0.01. n = 6 WT, n = 6 *Maf*^IECKO^. Scale bar, 50μm. **d**. Goblet cell hyperplasia upon loss of *Maf*. Quantification of number of goblet cells per villi and their diameter following periodic acid-Schiff staining. Scale bar, 50μm. (mean ± SD). *, p < 0.01. n=6 WT; n=6 *Maf*^IECKO^.

Tuft cells are rare chemosensory epithelial cells that play important roles in gut immune functions and intestinal regeneration (Howitt et al., 2016; Schneider et al., 2019; von Moltke et al., 2016; Westphalen et al., 2014). Tuft cell expansion has recently been reported to induce adaptive gut lengthening to preserve nutrient extraction capacity of the intestine (Schneider et al., 2018) and mucus-producing goblet cells have an important role in defence and repair of the small intestine mucosa (Renes et al. 2002). Staining for the tuft cell marker DCLK1, tuft/enteroendocrine cell marker PROX1, enteroendocrine marker CHGA or Paneth cell marker lysozyme (Cetin et al., 1989; Gerbe et al., 2009; Peeters and Vantrappen, 1975; Yan et al., 2017) revealed an expansion of tuft cells, but not enteroendocrine or Paneth cells in *Maf*^*ECKO*^ small intestine (**Fig.3c and Fig.S4c**,**d**). We also observed increased size but not number of goblet cells in the absence of *Maf* (**Fig.3d**) as determined by periodic acid – Schiff staining.

Collectively, these results show that loss of *Maf* in enterocytes promotes expansion or hyperplasia of selected secretory cell types that do not express c-MAF but promote intestinal regeneration and repair (May et al., 2014). While the exact nature of molecular signals involved requires further investigation, these results allow us to propose a model in which an altered enterocyte maturation program in the absence of *Maf* leads to tuft cell expansion and goblet cell hyperplasia, which in turn drives a compensatory enlargement of the small intestine, thus preserving a global metabolic balance of the organism.

### Epithelial *Maf* loss impedes recovery after acute intestinal injury

Since loss of *Maf* under steady state conditions resulted in intestinal adaptation, we next tested whether absence of epithelial *Maf* affects responses to acute intestinal injury. Administration of the potent chemotherapeutic agent methotrexate (MTX) depletes highly proliferative crypt progenitor cells, resulting in villus degeneration, transient malabsorption, and body weight loss (Sonis et al., 2001; Visentin et al., 2012). The intestinal epithelial lining is then rapidly regenerated from relatively quiescent columnar base stem cells leading to complete pre-treatment body weight recovery (Tian et al., 2011). WT and *Maf*^IECKO^ mice treated with MTX displayed a similar decrease in body weight until day 4 (**Fig.4a**). As expected (Aparicio-Domingo et al., 2015), the rapid weight recovery was observed in WT mice after the peak of damage. However, *Maf*^IECKO^ failed to gain weight leading to a tendency to decreased animal survival (**Fig.4a, Fig.S5a**).

**Figure 4.**
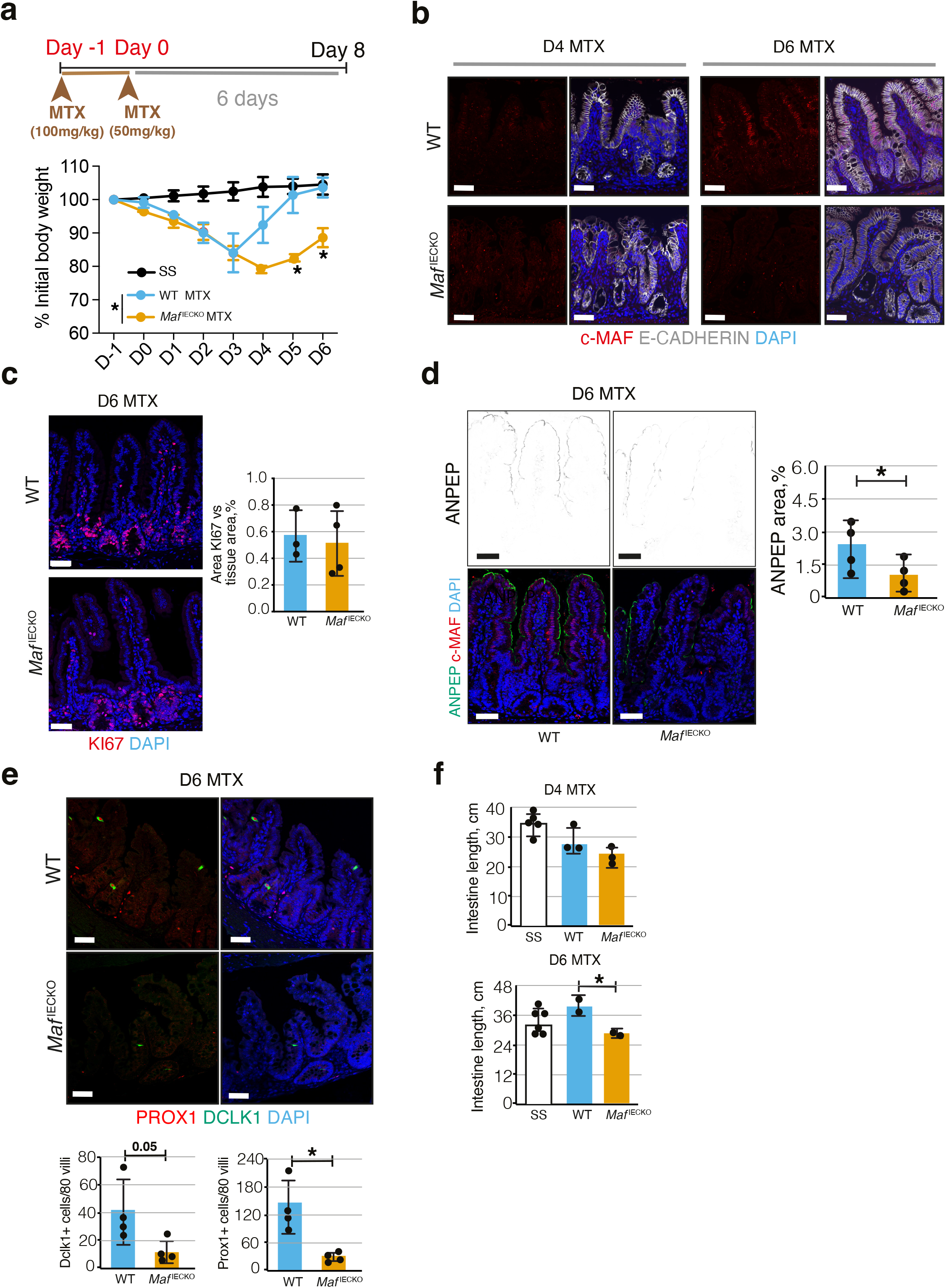
Defective expression of nutrient transporters and tuft cell expansion upon *Maf* loss impedes recovery after anti-metabolite treatment. **a**. Loss of *Ma*f prevents recovery after methotrexate (MTX) treatment. Percentage of initial body weight, mean ± SD. n.s.; *n* = 5 WT; *n* = 5 *Maf*^IECKO^ MTX mice treated and n= 4 WT and n= 4 *Maf*^IECKO^ control mice at the start of the treatment. **b**. c-MAF is downregulated at the peak of MTX treatment (day 4) and recovered at day 6 after treatment initiation. Staining for c-MAF (red), E-cadherin (white) and DAPI (blue). Scale bar, 50μm. **c**. No differences epithelial cell proliferation in *Maf*^IECKO^ mice. Staining for Ki67 (red) and DAPI (blue). Scale bar, 50μm. Mean ± SD. n.s. n=3 WT; n=3 *Maf*^IECKO^. **d**. ANPEP is reduced at day 6 after MTX upon *Maf* depletion. Staining for ANPEP (green), c-MAF (red) and DAPI (blue). Scale bar, 50μm. Percentage of area per tissue in WT and *Maf*^IECKO^ mice (mean ± SD). *, p < 0.05. n=3 WT; n=3 *Maf*^IECKO^. **e**. Tuft cells are reduced upon *Maf* loss at day 6 after MTX treatment. Staining for PROX1 (red), DLCK1 (green) and DAPI (blue). Number of PROX1^+^ and DCLK1+ cells in 80 villi. (mean ± SD). *, p < 0.05. n = 4 WT, n = 4 *Maf*^IECKO^. Scale bar, 50μm. **f**. Decreased length of small intestine in *Maf* depleted mice at day 6 after MTX treatment. Length of small intestine (cm) in WT and *Maf*^IECKO^ at day 4 and 6 after MTX initiation *, p < 0.05. n= 5 WT and n=6 *Maf*^IECKO^ control mice (PBS)-steady state (SS); n = 5 WT MTX treated, n = 5 *Maf*^IECKO^ MTX treated.

In WT mice, treatment with MTX induced rapid loss of epithelial c-MAF expression, which was regained during the regeneration phase (**Fig. 4b**). As MTX treatment targets proliferative cells, we analyzed epithelial cells during the recovery phase by staining for proliferation marker Ki67. In agreement with lack of expression of c-MAF in this compartment (**Fig.1b**), we did not observe differences in regenerating crypt proliferation or villus length between WT and *Maf*^IECKO^ mice (**Fig.4c**). The intestinal inflammatory response as assessed by expression of *Il6, Tnfa, Ifng, Il1b, Cd45* (**Fig.S5b)**, villi regeneration or overall maturation of the intestinal epithelium, as determined by KRT20 staining (**Fig.S5c)** were not significantly different in WT vs *Maf*^IECKO^ mice. However, levels of ANPEP (**Fig.4d**) were diminished at day 6 after MTX in *Maf*^IECKO^ mice as compared to WT animals. Furthermore, unlike in the steady state, we found that the intestines of MTX-treated *Maf*^IECKO^ harbored significantly fewer tuft but not goblet cells and the length of the small intestine was shorter as compared to WT mice (**Fig.4e,f** and data not shown). Collectively, these results indicate that *Maf* impairs the ability of mice to recover after acute intestinal epithelial damage due to a diminished/altered nutrient transport capacity of the regenerating differentiated epithelium and the inability to rapidly mount tuft cell-mediated compensatory intestinal lengthening.

In summary, our study sheds light on the fundamental novel role of c-MAF for maintaining a zonated transcriptional program of the differentiated small intestinal epithelium and its role in gut recovery after acute intestinal injury. It also highlights the importance of the cross talk between differentiated enterocytes and secretory intestinal cell populations to promote long term intestinal adaptation.

## Acknowledgements

We thank A. Moor for useful discussion, S. Nassiri for the initial analysis of RNA seq data, C. Beauverd for mouse genotyping, colony maintenance and immunostainings and J. Huelsken for providing mouse Rspo1. Mouse Pathology, Flow Cytometry, Animal, Cellular Imaging and Genomic Technology Facilities of the University of Lausanne are gratefully acknowledged. This work was supported by the Swiss National Science Foundation (CRSII5_177191, 31ER30_160674, 31003A-156266 and CRSK-3_190200 to T.V.P.,) and interdisciplinary grant of the Faculty of Biology and Medicine of UNIL (to T.V.P.).

## Author contributions

A.G.L. and T.V.P. planned the study and wrote the paper and T.V.P. supervised the research.

A.G.L. prepared the mouse models, performed the mouse studies, and analysed the data. T.W. contributed to figures 1a, e; S2b, c; 2d, e, g, h, S3b, c, d. O.M. helped on mice dissection for figures 2, 3. B.P.L provided samples for figure S1 and G.V. provided *Maf*^fl/fl^ mice. All authors read, commented on and approved the manuscript.

## Resource Availability

### Lead Contact

Further information and requests for resources and reagents should be directed to and will be fulfilled by the Lead Contact, Tatiana Petrova.

### Materials Availability

This study did not generate new unique reagents

## Data availabilty

Bulk RNA-seq raw sequence data and a table of counts per gene per sample reported in this paper have been deposited in the Gene Expression Omnibus (GEO) repository with the accession number GSE189852.

## Experimental models

### Mice

All experiments were approved by the Animal Ethics Committee of Vaud, Switzerland. *Maf* fl/fl mice (Giordano et al., 2015) were obtained from the laboratory of Gregory Verdeil. Mice were crossed with *Villin*-CreERT2 mice (el Marjou et al., 2004) to obtain *Maf*^fl/fl^;*Villin*-CreERT2 mice for *Maf* deletion in the intestinal epithelium. *Maf*^fl/fl^ mice were used as controls (WT). To activate the CreERT2 recombinase, 4-5 weeks old *Maf*^fl/fl^ and *Maf*^fl/fl^; *Villin*-CreERT2 received three intraperitoneal injections of tamoxifen (50 μg/g body weight) in Kolliphor EL (Sigma-Aldrich) every other day. For RNAseq analysis mice were sacrified one week after tamoxifen initiation. For all other experiments, mice were dissected after one-month deletion. For intestinal injury experiments, *Maf*^fl/fl^ and *Maf*^fl/fl^; *Villin*-CreERT2 were injected with 100 mg/kg of methotrexate (MTX) at day -1 and with 50 mg/kg at day 0 as described (Aparicio-Domingo et al., 2015). Body weight was monitored daily, and the intestine was collected at day 4 and day 6 after the last MTX injection. EdU (10mg/kg mouse; Invitrogen) was injected intraperitoneally 1 h or 48h prior to dissection.

## Methods details

### Staining procedures and image acquisition

*Paraffin sections*. 5μm sections were deparaffinized and subjected to heat-induced epitope retrieval using low or high pH Retrieval solutions (DAKO), incubated with antibodies and mounted in Fluoromount-G mounting medium supplemented with DAPI (eBioscience). The details of antibodies are provided in the **Reagents Table**. Alternatively, sections were stained using H&E (Harris modified hematoxylin and eosin solutions) (Sigma) and slides were mounted in Aquatex (Merck). Chromogenic detection was performed as previously described (Ragusa et al., 2014).

All images were taken using confocal Zeiss Zen v11.0.0.0 (Carl Zeiss) and analyzed using Imaris 8 (v.8.0.2) (Bitplane) and Photoshop CC v2015.1.2 (Adobe) softwares. Bright-field microscope images were taken using a Leica microscope MZ16 with a Leica Microsystems DFC 295 camera and LAS v4.2 software (Leica Microsystems).

### Image quantifications

For histological analyses at least 2-3 pictures per sample were taken with a Zeiss Zen v11.0.0.0 confocal microscope with 20x objective and quantified using ImageJ Fiji software. Keratin 20 (Krt20), ANPEP, DPP4 and APOA4, lysozyme-stained areas were normalized to total tissue area. For the quantification of epithelial cell proliferation, we measured the overlap of EdU/Ki67 signal over the E-cadherin selection. The images were transformed into 8-bit using ImageJ Fiji software and a threshold was set up. The same threshold was kept for all the pictures. The number of Prox1+, Dclk1+, ChgA+ cells was quantified manually in total of 60 villi in *Maf*^fl/fl^ and *Maf*^fl/fl^;*Villin*-CreERT2 mice. Goblet cell number per villi was manually quantified by following Periodic Acid Schiff (PAS) staining. Goblet cell diameter was analysed staining using particle analysis mask in Fiji. For quantification of crypt area, the ROI of crypts was quantified and measured. The pictures taken of every mouse were analyzed and the average value of those was plotted. GraphPad Prism 8 program was used to make the graphs and each dot represents a single mouse.

### Intestinal epithelial cells sorting

The second half of jejunum and the complete ileum were dissected and flushed with cold PBS. Peyer’s patches and mesenteric fat were removed, and the remaining small intestine was cut longitudinally and then washed with cold PBS. 0.5 to 1 cm pieces were incubated for 20 min in Dulbecco’s modified Eagle’s medium (DMEM) (Gibco) containing 10 mM EDTA with gentle stirring at 37°C. The tubes were vigorously shaken every 10 min to remove epithelial cells, intraepithelial lymphocytes, and mucus. The remaining intestinal tissues were incubated for 20 min at 37°C in digestion buffer containing DMEM, 2% serum, Liberase TL (0.2 mg/ml; Roche), de-oxyribonuclease (DNase) I (1 U/ml; Invitrogen), and 1% gentamicin with gentle stirring. To favour tissue dissociation, the pieces were mixed by pipetting up/down several times after 10- and 20-min. Isolated cells in the supernatant were harvested and kept on ice with a 1:1 volume of complete DMEM until the end of the digestion process. In total, three identical digestions were performed for the complete dissociation of the intestine. The cell suspension was passed through a 40-μm strainer, centrifuged for 6 min at 150*g* at 4°C, and resuspended in FACS buffer (PBS, 2% serum, and 2 mM EDTA). Cells were incubated with Fc block (antibody to CD16/32) for 20 min on ice and stained with conjugated antibodies in FACS buffer for 30 min. Following exclusion of dead cells using DAPI and gating on single cells, differentiated intestinal epithelial cells were selected as single Epcam+CD45-CD31-CD44-, gated and sorted directly in lysis buffer RLT containing *β-*mercaptoethanol (RNeasy micro kit, Qiagen). Cells were sorted on a BD FACSAria II (SORP) v8.0.1 cell sorter with BD FACSDiva software (BD Biosciences). Gating strategy is presented in **Figure S1**. Antibodies used for flow cytometry are listed in the **Reagents Table**.

### Tissue nucleic acid isolation and RT-qPCR

#### RNA isolation

Sorted cells, cultured organoids or total intestinal epithelium were transferred into RLT lysis buffer with *β-*mercaptoethanol (Qiagen) and snap-frozen on dry ice. RNA was isolated using Qiagen RNeasy Plus Micro Kit (Qiagen).

#### RT-qPCR

We used reverse transcriptor First Strand cDNA Synthesis Kit (Roche Diagnostics) and StepOnePlus Real-Time PCR instrument (Applied Biosystems) and SYBR Green PCR Master Mix (ThermoFisher) for qPCR analyses. Data were analyzed using the comparative Ct (ΔΔCt) method as described by the manufacturer. Gene expression was normalized to E-cadherin (**Fig.1d, Fig.S2, S5**) or 18S (**Fig.1f, Fig.S1**) levels. RT-qPCR results are shown as fold change over controls.

The nucleotide sequences of the real-time qPCR Primers used in this study are described in the **Reagents Table**.

### Bulk RNA-Seq

RNA QC, library preparations, and sequencing reactions were conducted at GENEWIZ, LLC. (South Plainfield, NJ, USA). Total RNA samples were quantified using Qubit 2.0 Fluorometer (Life Technologies, Carlsbad, CA, USA) and RNA integrity was checked with 4200 TapeStation (Agilent Technologies, Palo Alto, CA, USA). RNA sequencing library preparation used NEBNext Ultra RNA Library Prep Kit for Illumina by following the manufacturer’s recommendations (NEB, Ipswich, MA, USA). Briefly, enriched RNAs were fragmented for 15 minutes at 94 °C. First strand and second strand cDNA were subsequently synthesized. cDNA fragments were end-repaired and adenylated at 3’ends, and universal adapter was ligated to cDNA fragments, followed by index addition and library enrichment with limited cycle PCR. Sequencing libraries were validated on the Agilent TapeStation (Agilent Technologies, Palo Alto, CA, USA), and quantified by using Qubit 2.0 Fluorometer (Invitrogen, Carlsbad, CA) as well as by quantitative PCR (Applied Biosystems, Carlsbad, CA, USA). The sequencing libraries were clustered on two lanes of a flowcell. After clustering, the flowcell was loaded on the Illumina HiSeq instrument according to manufacturer’s instructions. The samples were sequenced using a 2×150 Paired End (PE) configuration. Image analysis and base calling were conducted by the HiSeq Control Software (HCS). Raw sequence data (.bcl files) generated from Illumina HiSeq was converted into fastq files and de-multiplexed using Illumina’s bcl2fastq 2.17 software. One mismatch was allowed for index sequence identification.

### Bulk RNA sequencing preprocessing and analysis

Sequencing reads were pseudoaligned using kallisto (v. 0.44.0) (Bray et al., 2016) against the *Mus musculus* reference transcriptome obtained from Ensembl (GRCm38.p6). Kallisto was run with default settings plus sequence bias correction.

Transcript level abundances quantified by kallisto were imported into R (v. 3.5.3). Conversion of Ensembl transcript identity to Ensembl gene identity, gene symbols, gene biotype and description was performed using biomaRt (v. 2.38.0) (Durinck et al., 2009). Counts for gene-level analysis were estimated using the “tximport” function of the tximport package (v. 1.10.0) (Soneson et al., 2015), with abundance estimates scaled using the average transcript length and library size (arguments set to: countsFromAbundance=“lengthScaledTPM” and dropInfReps=T). Next, we filtered out very lowly expressed genes by only retaining genes that were expressed at more than 0.45 counts per million in at least 2 samples. Differential gene expression analysis between c-MAF knockout and WT samples was performed using the “DESeq” function (with betaPrior=T) of the DESeq2 package (v. 1.14.1) (Love et al., 2014), and extracting the results using the function “results” with arguments addMLE=T and alpha=0.05. This yielded differential expression results for 15,558 genes, 6,751 of which had average normalized counts above 500.

Over-representation analysis of Gene Ontology (GO) terms (Ashburner et al., 2000; The Gene Ontology Consortium, 2019) was performed on genes with adj. p-value < 0.05 and average normalized counts above 500, separately for the up-regulated or down-regulated genes, using the “enrichGO” function of the clusterProfiler package (v.3.18.1 loaded in R v.4.0.2) (Yu et al., 2012) with argument minGSSize = 30.

### Expression level of *Maf* in publicly available data

We obtained microarray data of mouse tissues from GEO (GSE9954, (Thorrez et al., 2008). CEL files were imported into R (v. 3.5.3) using the “justRMA” function of the affy package (v. 1.60.0) (Gautier et al., 2004) and were converted to log_2_(normalized expression values). Annotation of microarray probes to gene symbols was obtained via the mouse4302.db package (v. 3.2.3) and we selected the probe for *Maf* with highest overall average expression (1456060_at) to create a heatmap using ComplexHeatmap (v. 1.20.0) (Gu et al., 2016).

We obtained single-cell RNA sequencing data of mouse intestinal epithelial cells from GEO (GSE99457) (Yan et al., 2017). The 3 output files of the cell ranger pipeline of all 10 samples were imported into R (v. 4.0.2) and pre-processed with Seurat (v. 4.0.4) (Hao et al., 2021). Counts were normalized with the logNormalize method and a scale factor of 10,000. A PCA was generated on the scaled values of the top 2000 variable genes selected with the variance-stabilizing method. A UMAP was generated using the first 20 PCs, followed by clustering of cells using the “FindNeighbors” and “FindClusters” functions with PCs 1 to 20 and a resolution parameter of 0.3. Clusters were annotated to cell types using known marker genes. To determine in which clusters the expression of *Maf* was highest, differential gene expression analysis among clusters was performed using the “FindAllMarkers” function with default parameters.

For the analysis of zonation, we obtained RNA sequencing data of mouse enterocytes from laser-micro-dissected villus sections from GEO (GSE109413, (Moor et al., 2018). Kallisto pseudo-alignment results were imported into R (v. 3.5.3) and pre-processed the same way as for the c-MAF KO and WT samples described above with tximport and biomaRt and normalized with DESeq2.

c-MAF expression in human and zebrafish intestinal epithelial cells was analysed using Single cell portal analysis tool (https://singlecell.broadinstitute.org/single_cell) and pre-publication dataset of human duodenum and ileum (https://singlecell.broadinstitute.org/single_cell/study/SCP817/comparison-of-ace2-andtmprss2-expression-in-human-duodenal-and-ileal-tissue-and-organoid-derived-epithelial-cells#study-summary) and dataset of intestine from zebrafish cultured under conventional conditions (GSE161855,Willms et al., 2021).

### Intestinal organoid culture

Intestinal stem cells were isolated from 12 weeks-old WT females and cultured as previously described (O’Rourke et al., 2016). For cell harvesting Matrigel was disrupted manually. For subsequent passaging, the organoids were dissociated by manual pipetting, the cells were washed, counted and 1000 cells/ 50 μl of Matrigel were cultured again as above. 48h after adding cell seeding, mouse recombinant BMP2 (100ng/ml, R&D) with or without 3μM DMH1 (Tocris) was added to the organoids and RNA collected 24h after.

### Statistical analysis

Statistical analysis was performed using Graphpad Prism 8 for MacOS (GraphPad). The number of mice analysed is indicated for each image in the figure legends. For *in vivo* experiments comparing WT and *Maf*^fl/fl^;*Villin*-CreERT2 animals, two-tailed unpaired Student’s t-test were performed to determine statistical significance between two means. For *in vivo* experiments comparing control and methotrexate-treated WT and *Maf*^fl/fl^;*Villin*-CreERT2 animals we used one-way ANOVA with Tukey post hoc test. Scattered dot plot data are shown as mean ± SD, where each single dot represents an individual mouse. p-value is stated in each figure. Differences were considered statistically significant at p < 0.05.

## Reagents Table

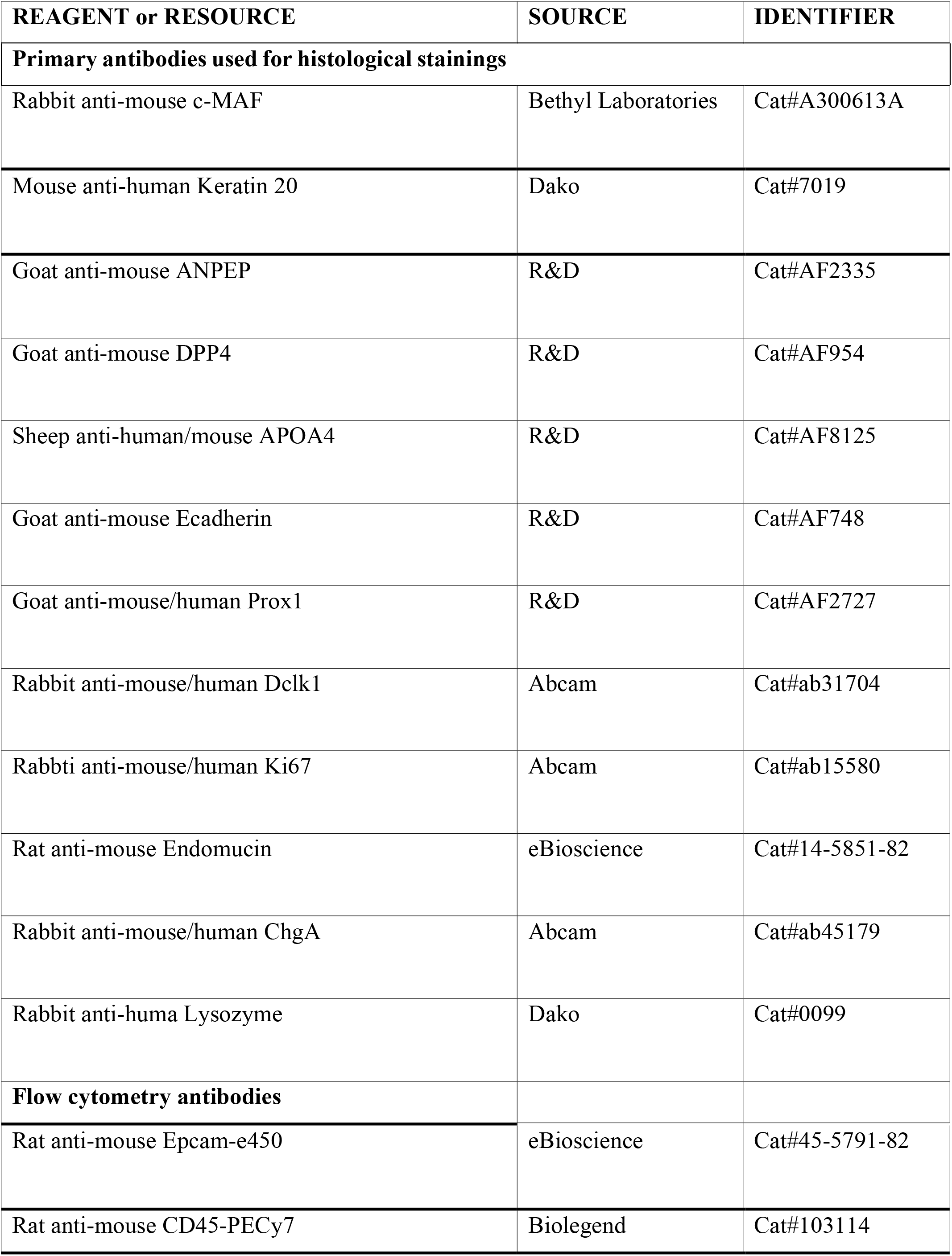
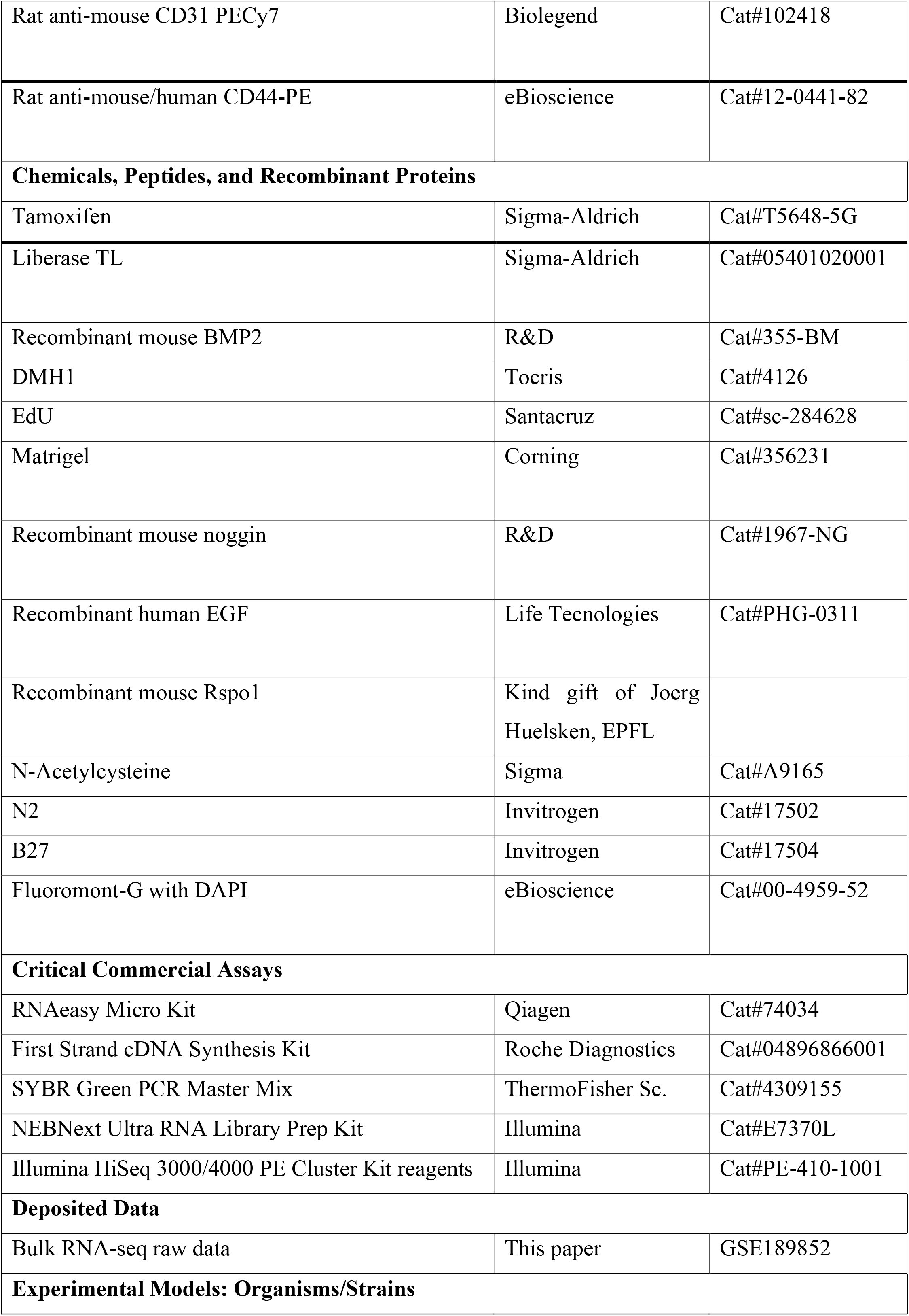
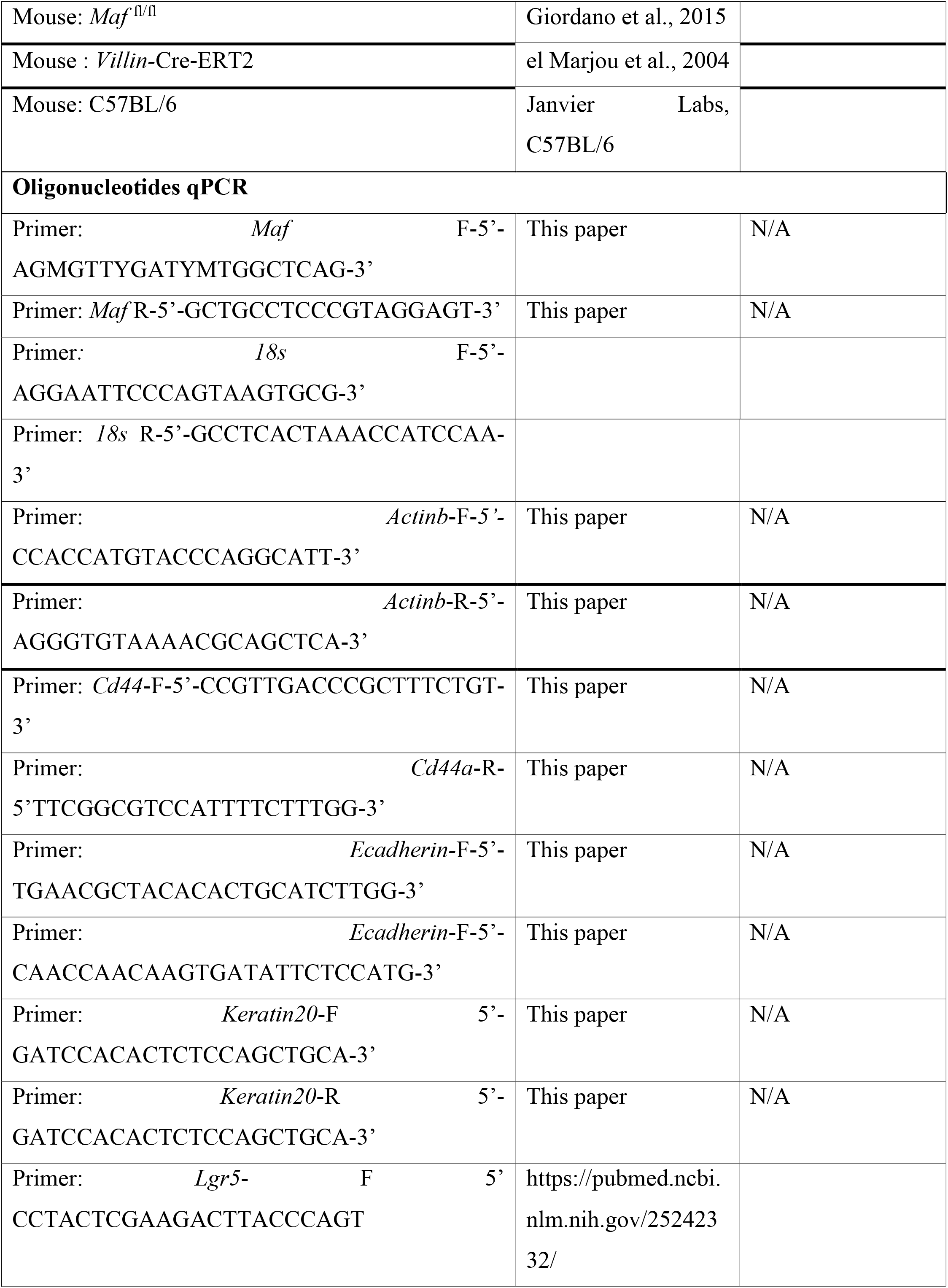
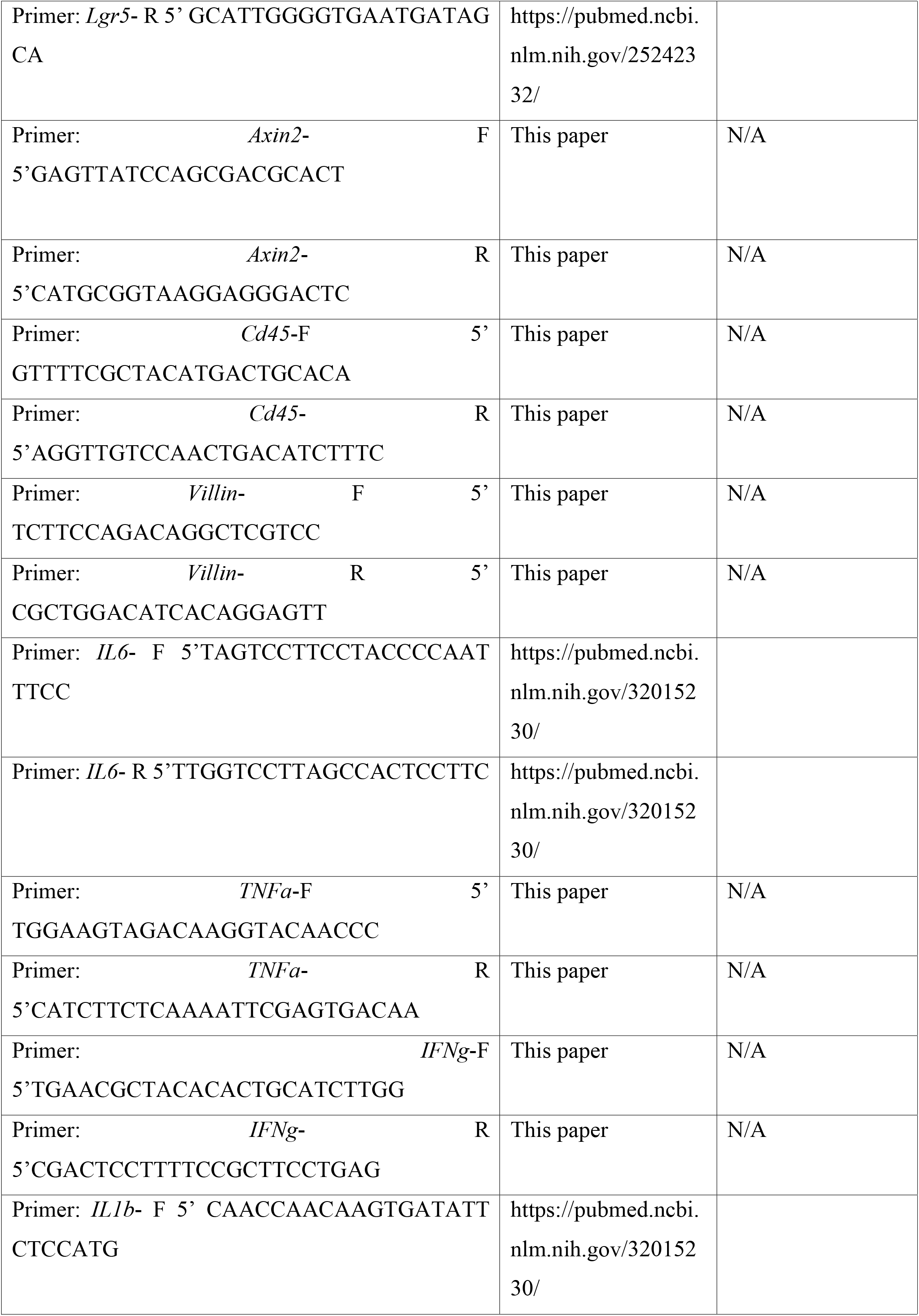
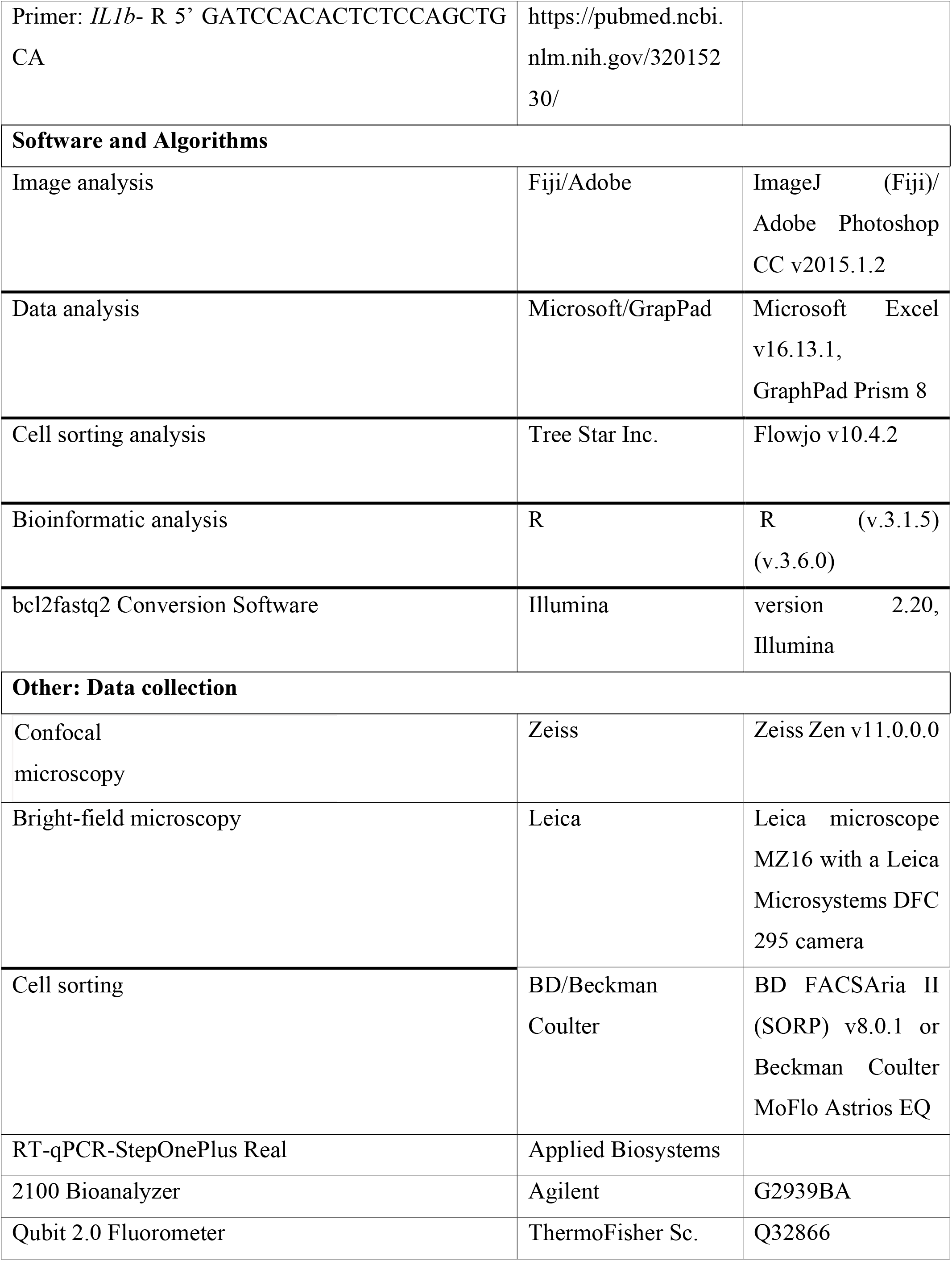

