## Supplementary figure and table legends for "c-MAF maintains the transcriptional program of enterocyte zonation and the balance of absorptive/intestinal secretory cell types"

### Supplementary figure legends

**Figure S1. c-MAF is expressed in adult mouse enterocytes**

**a.** Flow cytometry sorting strategy for isolation of enterocytes from 6-week-old mice. Sorted enterocytes are Epcam+ CD45-CD31- CD44-. **b**. Sorted CD44+ and CD44- intestinal epithelial cell purity analysis. mRNA levels of *Lgr5, Axin2, Cd45* and *Villin* normalized to 18S. **c.** UMAP plot of Yan et al., 2017 dataset. Thirteen major clusters of gut epithelial cells were identified. *Maf* expression per cell (ln[normalized counts + 1]) overlaid on UMAP plot. **d**. *Maf* is expressed in enterocyte clusters. Dot plot of markers in all clusters from Yan et al., 2017 dataset. The color code indicates scaled average expression level in each cluster, and the dot size denotes the percent of cells in each cluster expressing the given gene. Annotation of all clusters based on specific cell type markers (Yan et al., 2017).

**Figure S2. c-MAF expression in mouse intestine from development to adulthood**

**a.** Chromogenic staining for c-MAF along murine small intestine and colon. Scale bar, 50 µm. **b.** c-MAF is absent in the murine gut epithelium at E12.5. Staining for c-MAF (green), endomucin (red), PROX1 (white) and DAPI (blue). Scale bar, 200 µm. **c.** c-MAF is expressed in the murine gut epithelium at E17.5. Staining for c-MAF (green), E-cadherin (white), DAPI (blue). Scale bar, 50 µm. **d.** c-MAF is expressed in the murine gut at postnatal stage P4. Staining for c-MAF (green), E-cadherin (white), DAPI (blue). Scale bar, 50 µm. **e.** *Maf* mRNA relative levels in murine gut epithelium from postnatal day 8 to day 50. (mean ± SD). *, $p < 0.05$. N=4 WT mice.

**Figure S3. Aminoacid and bile acid transporter genes induced upon loss of *Maf* in adult intestine**

**a.** No compensation by other large or small Maf family members upon *Maf* depletion. *Mafb* and *Mafg* expression levels in Epcam+ CD44- RNA samples from WT and *Maf*$^{IECKO}$ mice. n.s. N=3 WT; n=2 *Maf*$^{IECKO}$ mice. **b.** Epithelial cell migration is unchanged in *Maf*$^{IECKO}$ mice. Staining for EdU (red) and DNA (blue). Scale bar, 50 µm. Quantification of EdU+ epithelial cells per length of villi. mean ± SD. n.s. $n = 3$ WT; $n = 3$ *Maf*$^{IECKO}$ 1h and 48h after EdU administration. **c.** Volcano plot of differentially expressed transcripts in *Maf*$^{IECKO}$ *versus* WT mice. Color scale indicates genes included in different GO pathways. **d**. *c-MAF* is highly expressed in differentiated human ileal enterocytes but not in progenitor/stem cells or secretory lineages. Dot plot of the indicated transcripts from human ileum dataset (https://singlecell.broadinstitute.org/single_cell/study/SCP817/comparison-of-ace2-and-tmprss2-expression-in-human-duodenal-and-ileal-tissue-and-organoid-derived-epithelial-cells#study-summary). The color code indicates the expression level in each cluster, and the dot size denotes the percent of cells in each cluster expressing the given gene. **e.** Zebrafish lysosome-rich enterocytes (LRE2) involved in protein catabolism and intestinal epithelial cells involved in nutrient absorption express high levels of *mafa*. Dot plot of indicated transcripts from GSE161855 (Willms et al., 2021). The color code indicates the expression level in each cluster, and the dot size denotes the percent of cells in each cluster expressing the given gene. EC: intestinal epithelial cell.

**Figure S4. Enteroendocrine cells remain unchanged upon *Maf* loss in enterocytes**

**a.** Colon length in *Maf*$^{IECKO}$ mice. Data shown as mean ± SD. Scale bar, 1 cm. n.s. *n* = 6 WT; *n* = 5 *Maf*$^{IECKO}$. **b.** No differences in the number of crypt fission events upon *Maf* loss. Staining for E-cadherin (red) and DAPI (blue). Scale bar, 50 µm. (mean ± SD). Quantification of number of double crypts per 0.1cm of tissue. n.s. *n* = 6 WT; *n* = 6 *Maf*$^{IECKO}$. **c.** ChgA+ cells remained unchanged upon *Maf* loss. Staining for ChgA (green) and DAPI (blue). Scale bar, 50 µm. Number of ChgA+ cells in 60 villi analyzed, mean ± SD. n.s. *n* = 3 WT; *n* = 4 *Maf*$^{IECKO}$. **d.** Loss of *Maf* does not alter Paneth cells. Staining for Lysozyme (green) and DAPI (blue). Scale bar, 50 µm. Lysozyme area in tissue area of WT and *Maf*$^{IECKO}$ mice, mean ± SD. n.s. *n* = 3 WT; *n* = 4 *Maf*$^{IECKO}$.

**Figure S5. *Maf* loss in enterocytes prevents recovery after anti-metabolite treatment**

**a.** Loss of *Maf* reduces survival of mice after methotrexate (MTX) treatment. Two consecutive doses of MTX were used and mice followed for a week. Mice percent survival; *n* = 5 WT; *n* = 6 *Maf*$^{IECKO}$ mice**.** **b.** Immune status is not altered upon *Maf* loss at day 6 after MTX. Relative mRNA levels for *Maf, Il6, Tnfa, Ifng, Il1b, Cd45* normalized to *Cdh1* expression. n=2 WT; *n* = 2 *Maf*$^{IECKO}$ mice. **c**. Keratin 20 levels do not change upon *Maf* loss after MTX treatment. Staining for c-MAF (red), K20 (green) and DAPI (blue). Scale bar, 50 µm.

### Supplementary tables

**Table S1.**

**A.** Differential gene expression between *Maf*$^{IECKO}$ vs WT (RNAseq) samples, obtained using DESeq2 (v. 1.14.1).

**B.** Gene sets significantly over-represented among the significantly up-regulated and down-regulated genes in *Maf*$^{IECKO}$ enterocytes versus WT mice (RNAseq)

**C.** Upregulated and downregulated genes in the villus zones described in (Moor et al., 2018) in *Maf*$^{IECKO}$ vesus WT (RNAseq) Villus region from (Moor et al., 2018), where each of the differentially expressed genes between *Maf*$^{IECKO}$ and WT is most highly expressed. The up- or down-regulated genes are sorted within each villus region according to adj. p-value of the comparison *Maf*$^{IECKO}$ vs WT. The villus region with the highest expression level was determined by averaging the expression level of each gene across the 3 replicates included in Moor et al., 2018.
