## Supplementary figures and images for "c-MAF maintains the transcriptional program of enterocyte zonation and the balance of absorptive/intestinal secretory cell types"

### Supplementary figure 1

#### Supplementary figure 1

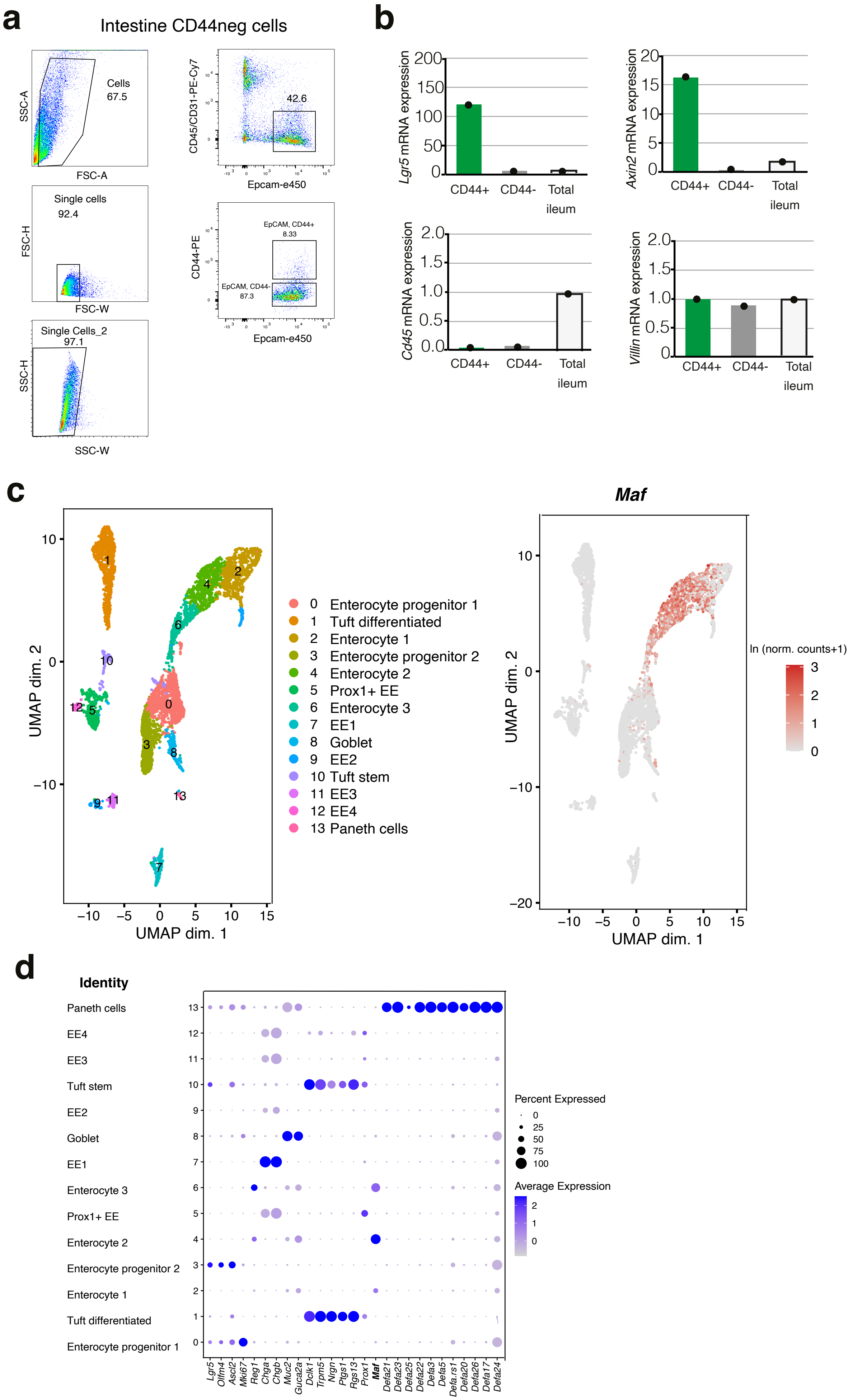

### Supplementary figure 2

#### Supplementary figure 2

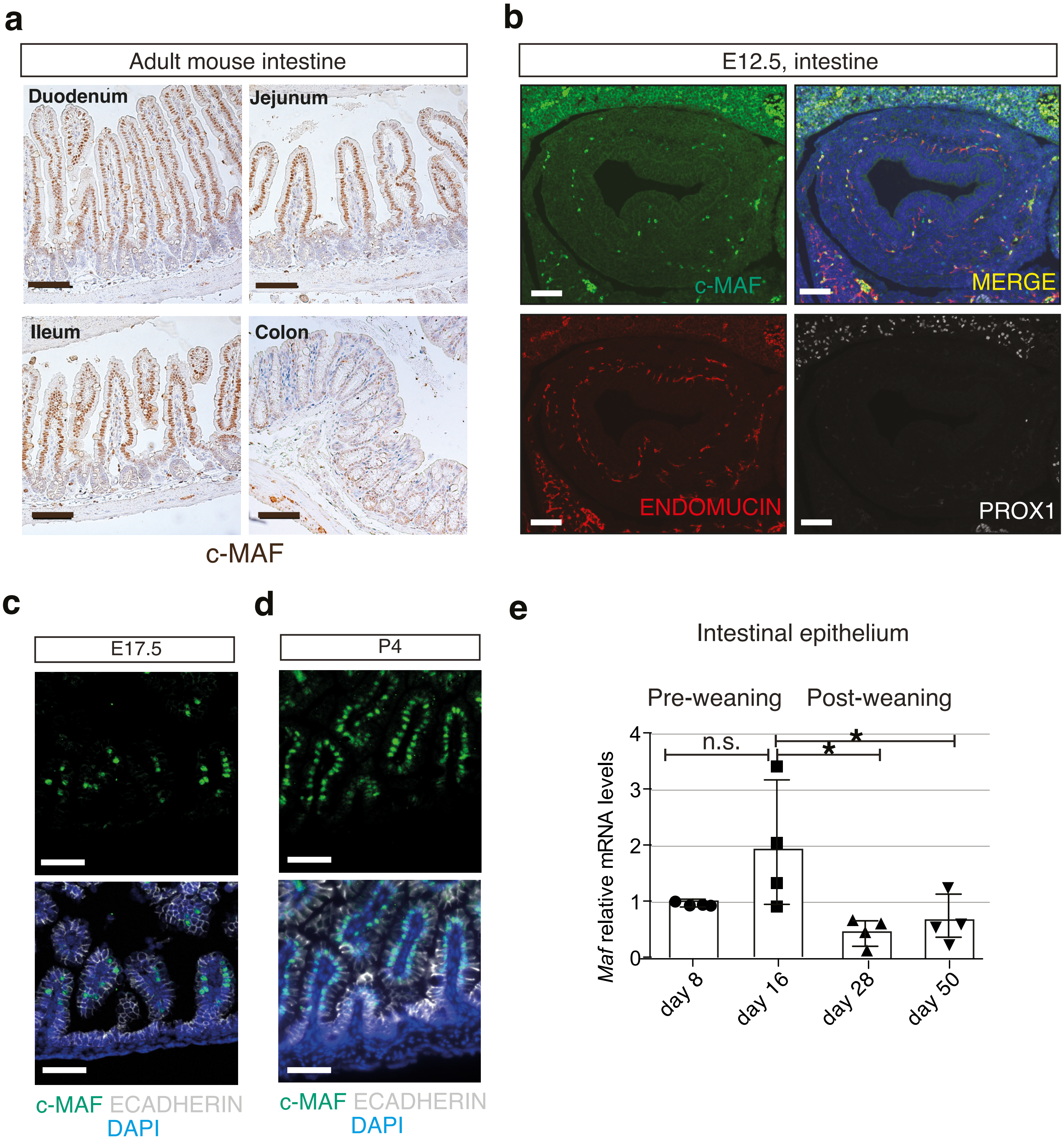

### Supplementary figure 3

#### Supplementary figure 3

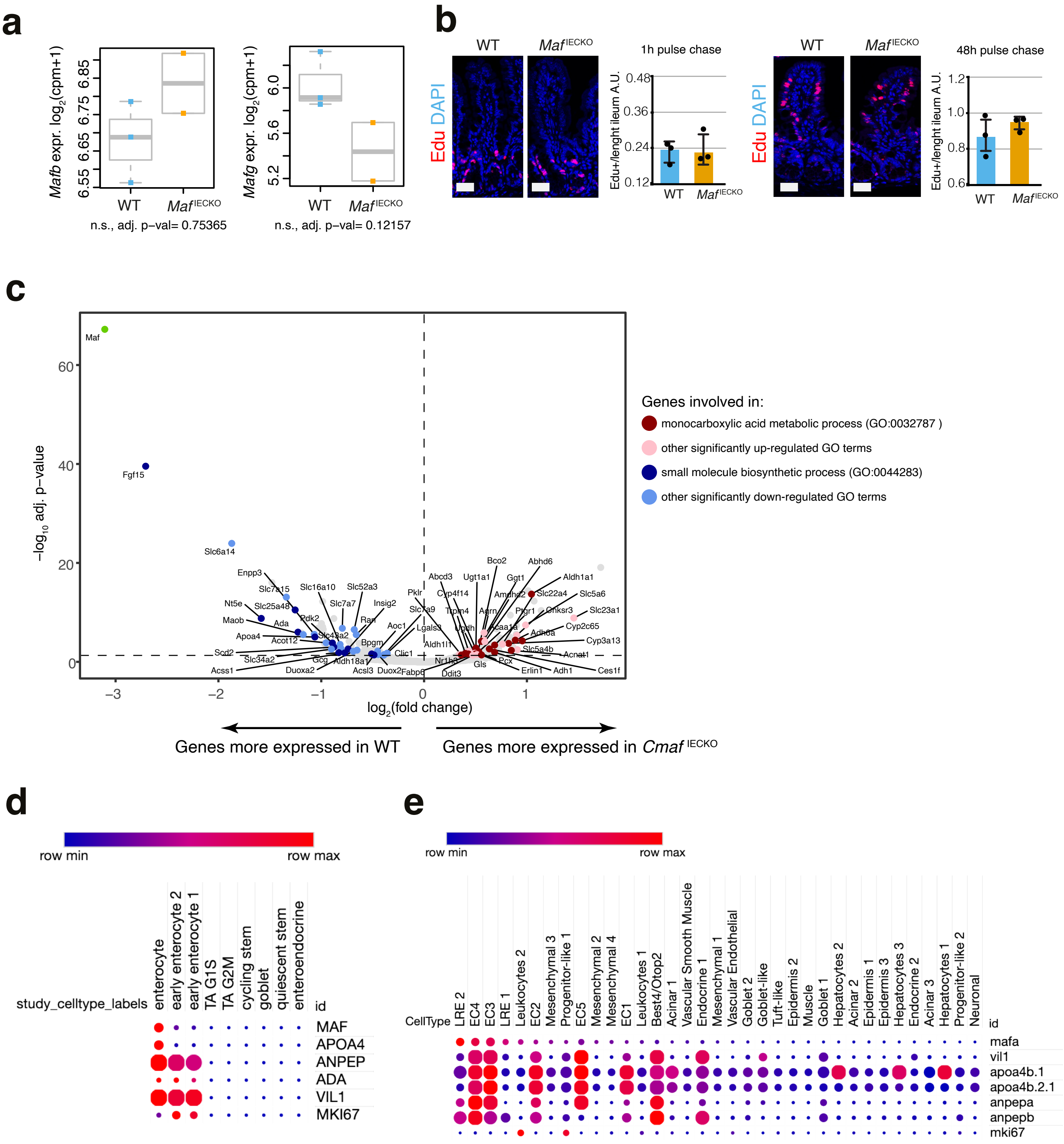

### Supplementary figure 4

#### Supplementary figure 4

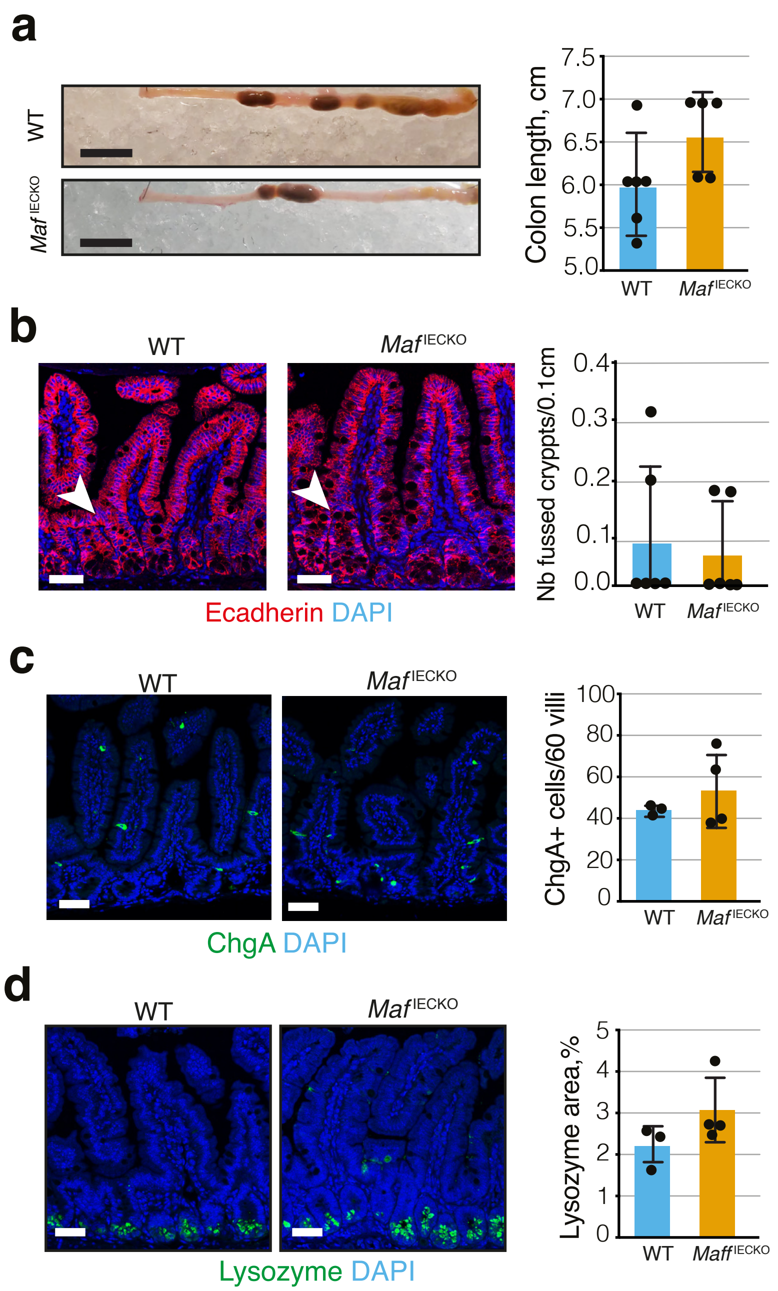

### Supplementary figure 5

#### Supplementary figure 5

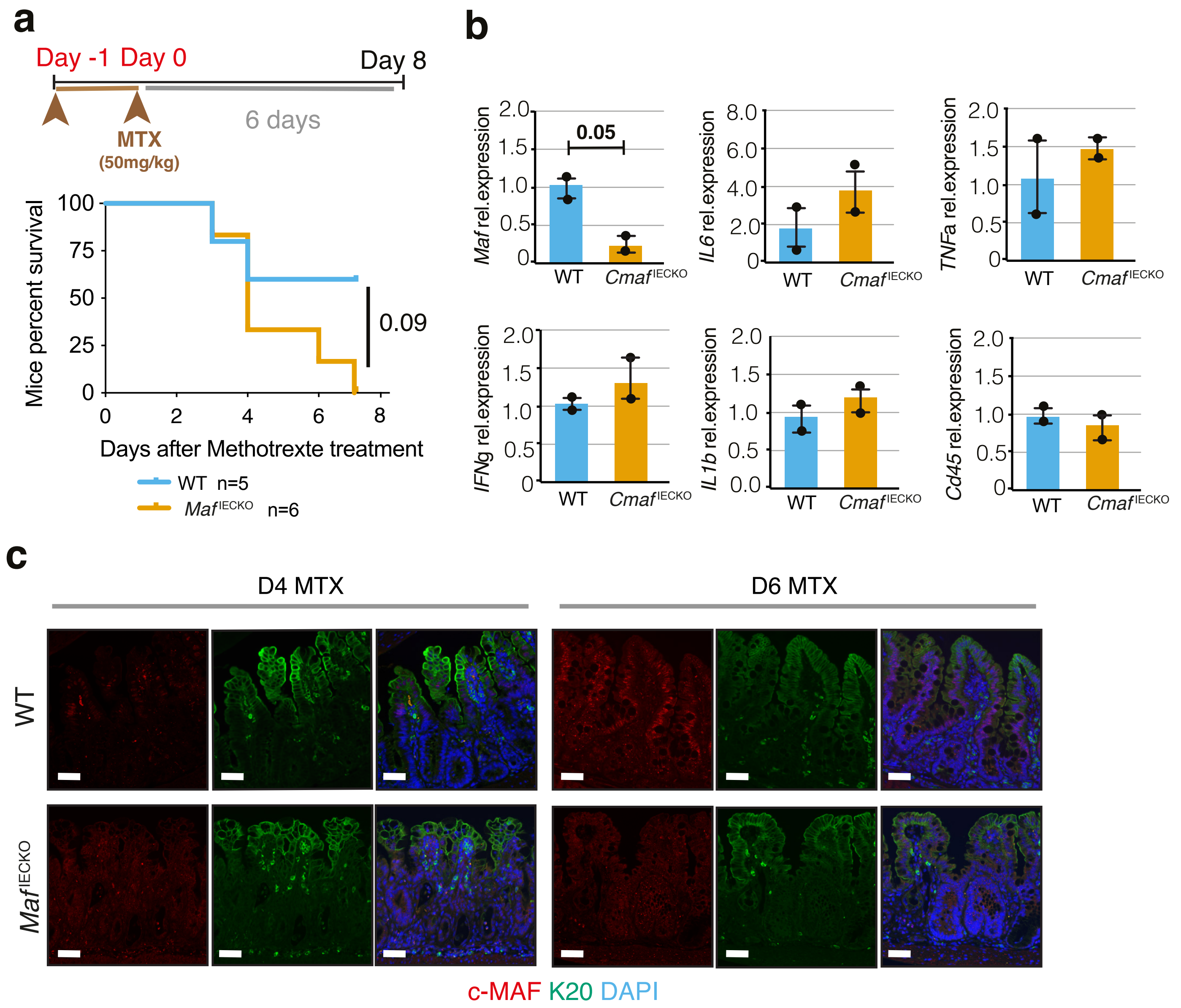
